# Deep Learning of Ligand-bound RNA Tertiary Structures Diverges from Learning Unbound Ones: A Case Study Using The gRNAde Software

**DOI:** 10.1101/2023.09.13.557627

**Authors:** Tajveer S. Dhesi, Alyssa W. Bannister, Amirhossein Manzourolajdad

## Abstract

Modeling the relationship between the native RNA sequence and its *in-vivo* structure is challenging, partly due to the flexible nature of the RNA molecular structure. In addition, the RNA structure can take on different conformations in the presence of specific molecules, metabolites, temperatures or other signaling and environmental factors, making it difficult to construct a universal statistical model for the sequence-structure relationship of the RNA. Using a Geometric-Vector-Perceptron Graph Neural Network architecture, Joshi, et al. predict the RNA sequence from its given 3D structure with good performance and on a dataset including RNA structures of different type and length, namely RNAsolo. In this work, using the Authors open-source software package, gRNAde, we confirm their results on a more updated version of RNAsolo and for structure of different resolution, confirming the ability of the algorithm to capture RNA structural features and generalize to sequences of different lengths. We did observe, however, that performance on riboswitches is lower than expected that RNAs whose structure has been resolved while being bound to a ligand, such as riboswitches, may require a statistical model that diverges from those of native structures.

## Introduction

Modeling all the governing forces that act upon the RNA molecular structure simultaneously can be challenging. The RNA molecule has a flexible structure (Hyeon, Dima et al. 2006), which can take on different conformations in the presence of specific molecules, metabolites, temperatures or other signaling and environmental factors (Serganov and Nudler 2013). In addition, due to co-transcriptional folding, i.e., RNA folding at it is being transcribed, transcription speed and environmental factors can alter the final structure that an RNA stabilized into (Watters, Strobel et al. 2016). Although certain RNAs such as micro RNAs and ribosomal RNAs are evolved into folding into a single deterministic structure, certain others such as riboswitches perform their regulatory roles by being flexible enough to fold into two distinct conformations, one unbound and the other bound to a specific external factor, usually a ligand (Serganov and Nudler 2013).

Finding the relationship between the RNA molecular structure and its native sequence composition has been one of the main challenges of RNA-related bioinformatics studies. In some cases, the RNA is reduced to a two-dimensional schematic known as the secondary structure, whereas in other more comprehensive prediction tasks the complete 3D structure of the RNA, known as tertiary structure, is to be predicted. The ground truth of tertiary structures can be derived from experiments such as X-ray crystallography or Cryogenic electron microscopy (cryoEM). Similar to protein 3D structures, RNA structures are represented by the three-dimensional coordinates of each of their atoms and in the format of a Protein-Database (PDB) file.

In recent years, advances in deep learning such as those in Geometric Graph Neural Networks (Geomteric GNNs) done by (Bronstein, Bruna et al. 2021) have greatly contributed to prediction of geometrical structures in general and structures of biomolecules such as small molecules, proteins, and RNAs in specific. A recent work uses geometric GNNs to learn the tree-dimensional structure of the RNA molecule form its sequence (Townshend, Eismann et al. 2021) outperforming current methods. Graph neural networks have also been used in the reverse design application for both proteins and RNAs where the structure of the molecule is given but the sequences are missing. (Ingraham, Garg et al. 2019) apply GNNs to protein structures to predict amino-acid sequences of the protein. The approach predicts each amino acid of the protein sequence, separately. Prediction is based on previous amino acids as well as those that are spatially in proximity of the amino acid to be predicted. In a computational RNA design approach (Joshi, Jamasb et al. 2023), use the Geometric-Vector-Perceptron GNN (GVP-GNN) architecture presented in (Jing, Eismann et al. 2021) to predict RNA sequences from structures. Authors generalize to incorporating not only the structure and sequence information of one instance of structure but also those for other instances of structures derived from multiple experiments. Hence, for each sequence, multiple structures are averaged and then used in the learning as opposed to only one structure per sequence. The above approach, known as multi-state RNA design, was tested on a large-scale dataset, RNAsolo (Adamczyk, Antczak et al. 2022), and was shown to outperform ‘single state’ design, that uses only one structure per sequence, in improving native sequence recovery.

RNAsolo is a recent repository of RNA 3D structures. Structures are extracted from various sources including protein-RNA complexes and DNA-RNA hybrids. RNAsolo consists of all RNA structures including those with a single structure such as miRNAs and ribosomal RNAs, as well as those evolved to have alternative structures such as riboswitches. In this work, we use the gRNAde software to assess its performance specifically on riboswitches as opposed to all RNA molecules. The goal is to assess whether the performance of the multi-state RNA design on riboswitches follows other classes of RNA and to have a better understanding of the complexity of modeling the structure of different types of RNA molecules.

### Data Collection

Structures were gathered from RNAsolo (Adamczyk, Antczak et al. 2022) a recent repository of RNA 3D structures that stores available in RNA tertiary structures PDB format. The website is routinely updated and each month, more data is added to the repository. Four distinct datasets of structures, all downloaded from RNAsolo, are considered in this study. First dataset refers to the one used by (Joshi, Jamasb et al. 2023). It consisted of 11,538 structures all at resolution ≤3A° and referred to here by 3Aall. The second dataset was downloaded on August 10, 2023. We chose all structures at resolution ≤4A°, which resulted in a total of 13682 structures. We refer to this dataset as 4Aall. To retrieve all riboswitch sequences, our third dataset, we selected sequences having with the term ‘riboswitch’ and length between 10nt and 500nt from RNAsolo, which resulted in a total of 533 structures. We refer to this dataset as “switch”. Finally, as forth set of structures, we selected “ribosome” as a keyword, which resulted in a total of around four thousand sequences. Around 550 of these PDB files were randomly selected and the resulting dataset was called “unitary”, representing RNA molecules with a single deterministic conformation. Table 1 summarizes the statistics of the four datasets used in this work.

**Table 1.** TEST recovery for Multi-State RNA Design. RMSD Split was used to split data into training, validation, and test sets. Analysis was identical to that of (Joshi, Jamasb et al. 2023), which also describes computing Mean and Median sequence recovery. Parameter *k*, number of sampled structures, was set to 3. Number of epochs was set to 100. 4Aall refers the data downloaded on August 10, 2023, with structures at resolution ≤4A°. 3Aall refers to results reported by Joshi et. Al on the data downloaded at resolution ≤3A°. *Switch* refers to dataset of riboswitches as described in Methods. *Unitary* refers to dataset of sampled ribosome RNAs as described in Methods.

| Dataset | Unique Seqs. | No. of structures | Mean | Median |
| --- | --- | --- | --- | --- |
| 3Aall. (Joshi et al) | 4,707 | 11,538 | 62.02% $\pm 1.19$ | 65.03% |
| 4Aall | 3,764 | 13,682 | 58.00% $\pm 2.20$ | 61.41% |
| switch | 273 | 533 | 48.30% $\pm 1.546$ | 46.03% |
| unitary | 202 | 550 | 69.30% $\pm 1.69$ | 69.02% |

## Results

Statistics for all structures downloaded from RNAsolo, 4Aall, is shown in Fig. 1. The figure illustrates distributions of sequences length (Fig. 1A), number of structures per sequence (Fig. 1B), structural diversity, average pairwise root-mean-square distance (RMSD), per sequence (Fig. 1C), as well as bivariate distribution of sequence length and RMSD (Fig. 1D). Statistics of 4Aall is very similar to the statistics of 3Aall described in previous work (Joshi, Jamasb et al. 2023), with average sequence length being around 100nt and average RMSD of roughly less than 1.

**Figure 1.**
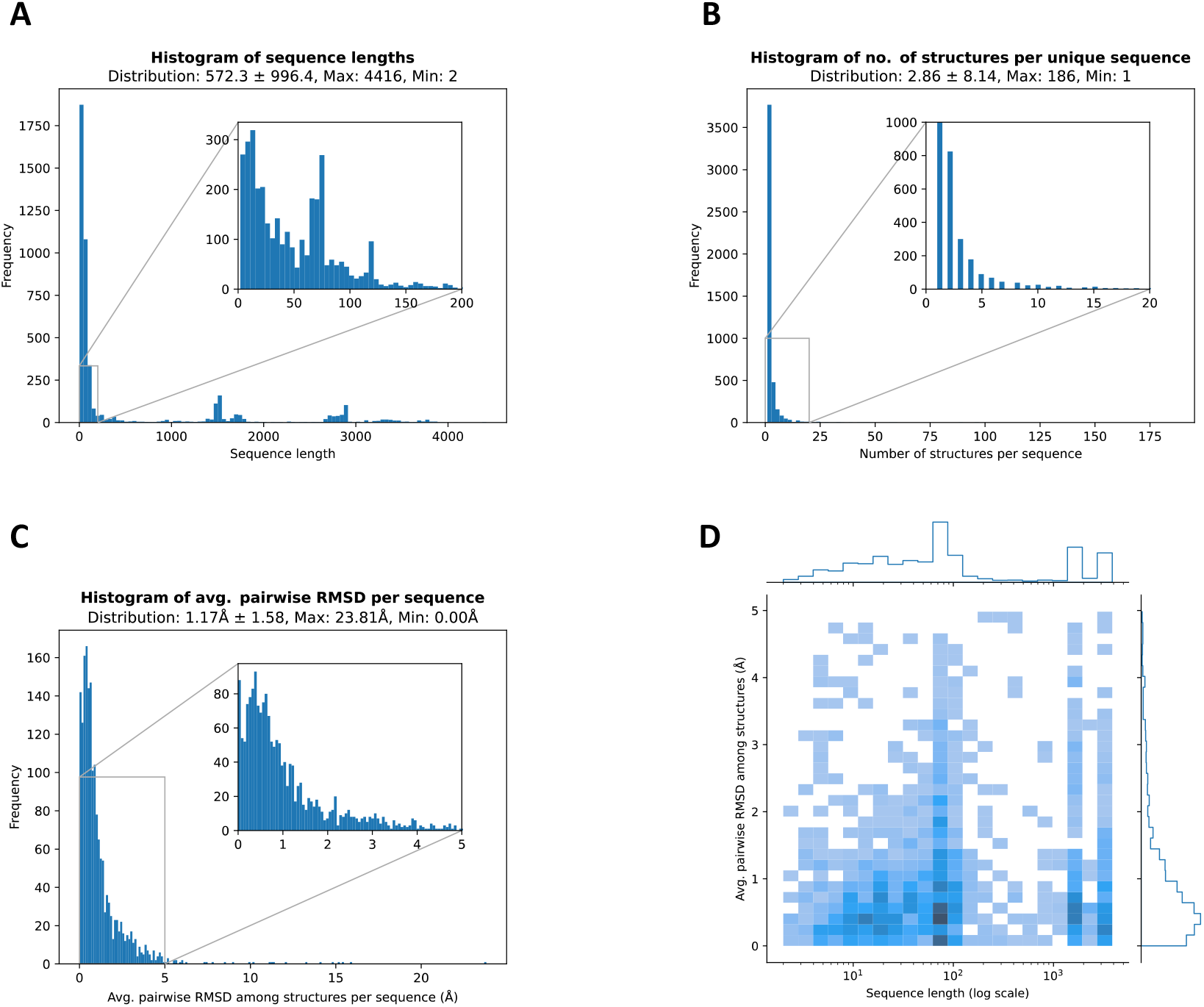
Data statistics for the 13682 structures at resolution ≤4A° downloaded from RNAsolo. The dataset is referred to as 4Aall. (A) shows histogram of sequence length. (B) illustrates number of structures available for each sequence. (C) depicts histogram of average pairwise Root-Mean-Square-Distance (RMSD) per sequence. (D) is a heatmap of sequence length and average RMSD.

Statistics for riboswitches, switch, is shown in Fig. 2. As we can see, although the number of structures in switch were much less than total of RNA structures, 533 vs 13,682 structures, their features such as sequence length and RMSD seem typical of total number of sequences (comparing Fig. 1 and Fig. 2).

**Figure 2.**
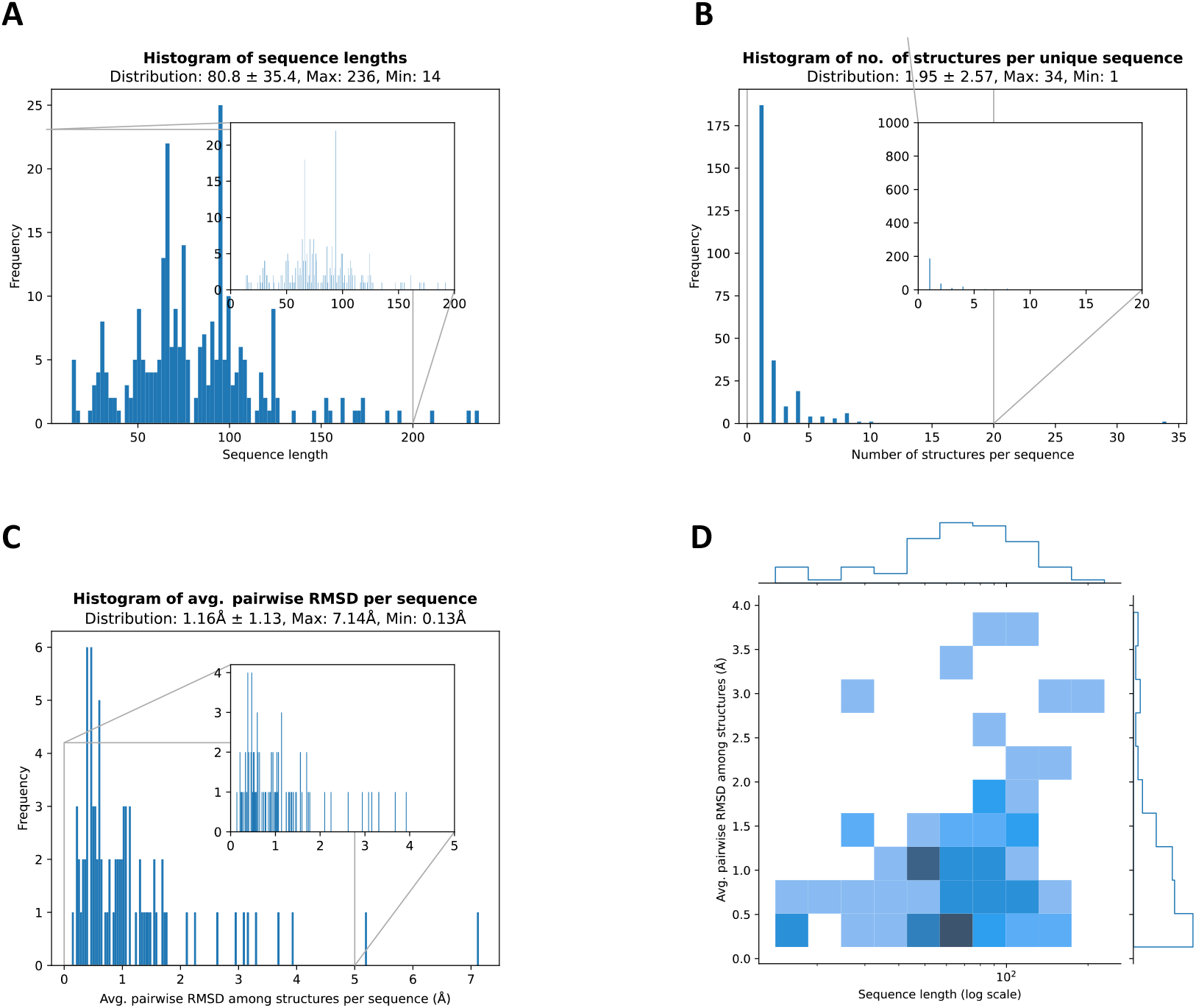
Data statistics for the 533 structures of the *switch* dataset. Sequences having term ‘riboswitch’ and length between 10nt and 500nt were downloaded from RNAsolo. (A) shows histogram of sequence length. (B) illustrates number of structures available for each sequence. (C) depicts histogram of average pairwise Root-Mean-Square-Distance (RMSD) per sequence. (D) is a heatmap of sequence length and average RMSD.

Statistics for *unitary* dataset is shown in Fig. 3. As we can see, although the number of structures of this dataset is similar to the switch dataset, both being around 500 structures, sequence lengths are very dissimilar. In fact, sequence length distribution of the *unitary* dataset follows a bimodal distribution that is centered around both 100nt and 1500nt. Overall, the switch dataset is more typical of the total number of structures than the *unitary* dataset.

**Figure 3.**
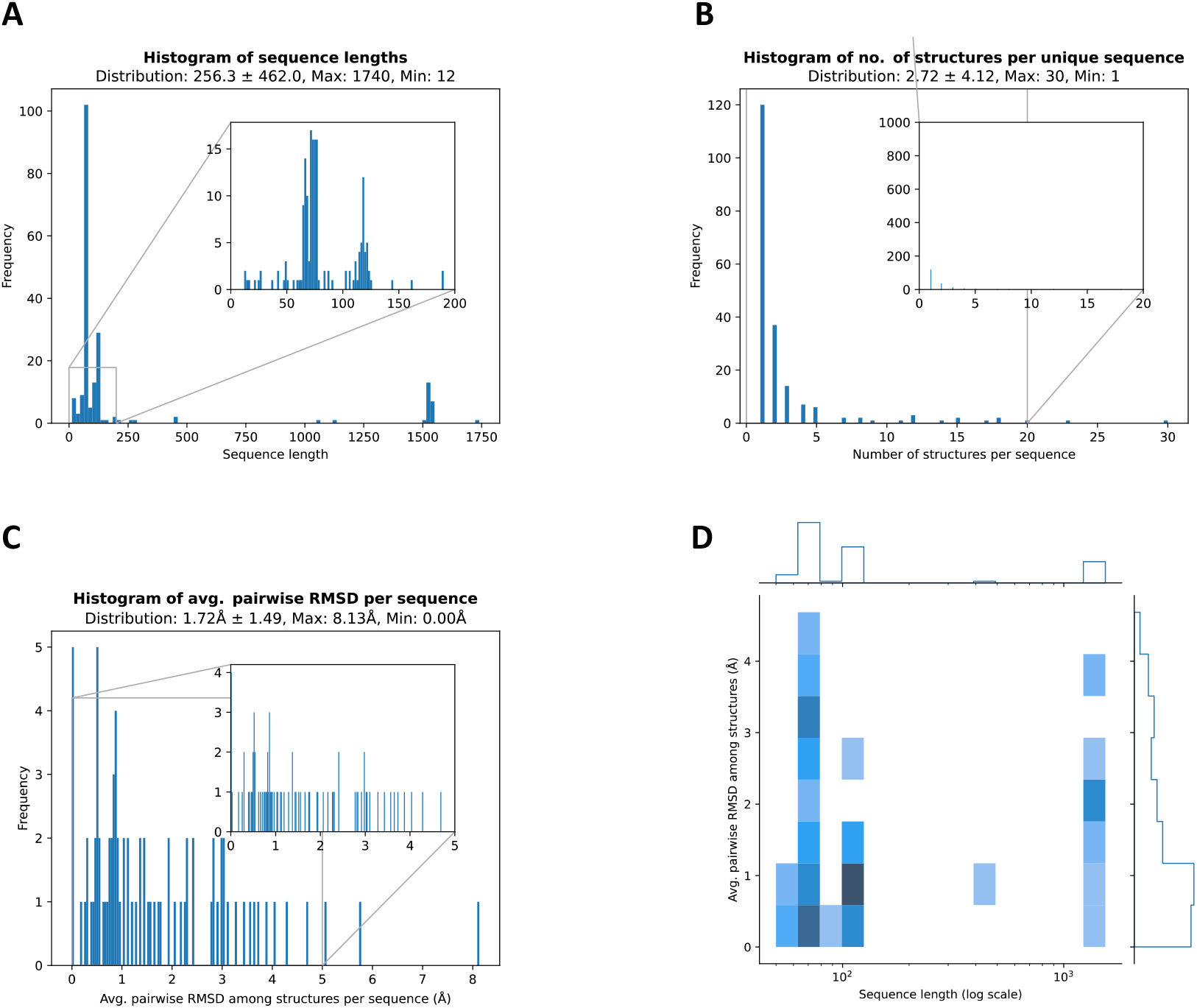
Data statistics for the 550 structures of the *unitary* dataset. Sequences having term ‘ribosome were downloaded from RNA solo. (A) shows histogram of sequence length. (B) illustrates number of structures available for each sequence. (C) depicts histogram of average pairwise Root-Mean-Square-Distance (RMSD) per sequence. (D) is a heatmap of sequence length and average RMSD.

The gRNAde software developed by (Joshi, Jamasb et al. 2023) was run on the three datasets. Performance here is defined as mean and median of test-set sequence recovery. Details of definition of sequence recovery as well as criteria for splitting sequences in to training, validation, and test sets, are all described in (Joshi, Jamasb et al. 2023). The procedure was applied in this work accordingly, except for when we needed to reduce the size of the test sets for *switch* and *unitary* datasets, due to smaller size of structure in these datasets. Instead of defining the size of the test set to 200 as done in (Joshi, Jamasb et al. 2023), we set the corresponding parameter to 20 for both *switch* and *unitary* datasets.

Table 1 describes performance of the multi-state design deep learning software, gRNAde, on all four datasets. As we can see, structures at resolution ≤4A° have a similar performance with those at ≤4 A° (Comparing 3Aall to 4Aall in Table 1). Performance on the 3Aall dataset, however, is slightly higher than the 4Aall dataset (medians 65.03% vs 61.41%). This slight difference in performance might be due to the latter having lower resolution structures.

The two datasets *switch* and *unitary* both have similar numbers of structures, 533 and 550, respectively as well as similar numbers of unique sequences within each set, 273 and 202, respectively. The performance of gRNAde on the switch dataset, however, is much lower than that of the *unitary* dataset (Comparing Medians 46.03% and 69.02%, Table 1). In fact, the performance on the *unitary* dataset is higher than that of all RNAs (Comparing medians 69.02% to 61.41%, Table 1). Interestingly, while sequence lengths within the switch dataset are much more similar to that of all RNAs (Comparing Fig. 2 to Fig. 1), the performance of gRNAde on the switch dataset is very different and much lower than that of all RNAs (Comparing 46.03% to 61.41%, Table 1).

The performance of gRNAde on *switch* and *unitary* was also evaluated under the parameter *k*. Parameter *k* is the number of sampled structures for each sequence during the training phase of deep learning. Default value for sampling structures is *k* = 3. It has been shown in table 1 of (Joshi, Jamasb et al. 2023) that the performance of *k* = 3 is superior to that of *k* = 1 (single-state baseline) regardless of data splitting strategies used for divide structures into training, validation, and test sets. In this work, similar to (Joshi, Jamasb et al. 2023), performance increases for the unitary dataset as we increase *k* from 1 to 3 (Comparing Medians 63.52% and 69.02%, Table 2). In stark contrast to results from all other three datasets, results for the switch dataset tend to have an opposite trend, with performance corresponding to *k* = 1 being either higher or similar to that of *k* = 3 (Comparing Means 50.00% ±1.485 and 48.30% ±1.546 as well as Medians 49.27% and 46.03%, Table 2).

**Table 2.** TEST recovery for Multi-State RNA Design. RMSD Split was used to split data into training, validation, and test sets. Analysis was identical to that of (Joshi, Jamasb et al. 2023), which also describes computing Mean and Median sequence recovery. Parameter *k*, number of sampled structures, was set to 1 and then 3. Number of epochs was set to 100. 4Aall refers the data downloaded on August 10, 2023, with structures at resolution ≤4A°. 3Aall refers to results reported by Joshi et. Al on the data downloaded at resolution ≤3A°. *Switch* refers to dataset of riboswitches as described in Methods. *Unitary* refers to dataset of sampled ribosome RNAs as described in Methods.

| | $k = 1$ | | $k = 3$ | |
| --- | --- | --- | --- | --- |
| Dataset | Mean | Median | Mean | Median |
| switch | 50.00% $\pm 1.485$ | 49.27% | 48.30% $\pm 1.546$ | 46.03% |
| unitary | 63.49% $\pm 1.108$ | 63.52% | 69.30% $\pm 1.69$ | 69.02% |

## Discussion and Conclusions

In this work, we evaluated the performance of the deep-learning-based multi-state RNA design, gRNAde, on different classes of RNAs with the focus on riboswitches. The algorithm was based on the application of Geometric GNNs to predict the nucleotides of an RNA sequence from its structure. The approach was designed and implemented by (Joshi, Jamasb et al. 2023). Using the gRNAde software our results are mostly similar to those presented in (Joshi, Jamasb et al. 2023). Using a more updated set of structures from RNAsolo, it was shown that the performance of gRNAde follows the same trend as for previous datasets (Table 1). Furthermore, performance on a smaller dataset of only size 550 which consisted of sparse distribution of sequence lengths (Fig. 3) also led to similar results, confirming the algorithm’s ability to capture RNA structural features and generalize to sequences of different lengths.

Results for riboswitches, however, different from what was expected. As we can see from Table 1, performance on the switch dataset is much lower compared to other sets of RNAs (Table 1). This poor performance is not necessarily due to non-structural features of our dataset. Comparing dataset statistics, Figs. 1, 2, and 3, shows that in fact, riboswitches have a similar length distribution to the complete set of RNAs, 4Aall. On the other hand, length distribution of the unitary dataset seems to be an outlier. Nevertheless, performance on the unitary dataset is higher than all other datasets (Table 1). Other factors such as number of unique sequences or structural diversity measures are very similar between *switch* and *unitary*, hence, neither sequence nor structural discrepancies are large enough to account for difference in performance. We investigated some of the riboswitches in RNAsolo and it seems that structures of most these riboswitches were resolved when the riboswitch was bound to a ligand or a molecule. Although the aptamers of riboswitches have been evolved for millions of years to be structurally conserved, it seems that the structures used here are not capturing the innate structure but the bound structure of the aptamer. We suspect that the use of ligand-bound structures might explain the low performance. The governing molecular forces for these bound RNA structures are not only different from those for innate structures but even different from each other, since each riboswitch binds to a different ligand, or metabolite than the other one. Hence, making it harder to derive a universal geometric model for all RNAs.

The performance of gRNAde on the two small datasets *switch* and *unitary* followed opposing trends once changing the number of sampled structures or parameter *k*. Default value for sampling structures is *k* = 3 which outperforms models with *k* = 1. This increase in performance is clearly seen on the unitary dataset where increasing *k* from 1 to 3 increases model performance (Comparing Medians 63.52% and 69.02%, Table 2). We observed, however, that in the *switch* dataset, model with *k* = 1 already has a higher performance than that with *k* = 3 (Comparing Means 50.00% ±1.485 and 48.30% ±1.546 as well as Medians 49.27% and 46.03%, Table 2). It is not clear why performance does not increase for riboswitches as parameter *k* increases. First, it may just be that we do not have enough data or have not repeated the test enough number of times to infer a lower performance for *k* = 3. Data splitting strategies may also partly explain this discrepancy in performance. Another speculation is that RNA molecules have been evolved to have sequences meticulously arranged to accommodate two overlapping structures. Predicting sequence of a riboswitch from only one of its structures, here the bound structure, may only partly recover the sequence of the riboswitch, given that there are less constraints for an RNA reverse design than the application of both unbound and bound structures.

Finally, RNA computational design might be facing yet another challenge once more data on riboswitches contain mutually exclusive bound and unbound conformations of the same sequence. Currently, the multi-state design algorithm presented in (Joshi, Jamasb et al. 2023) averages different structures before feeding them into the GNN (Equation 3 of (Joshi, Jamasb et al. 2023). The approach works naturally for cases where there are multiple experiments all resolving an identical structure, hence reducing noise. For the case of riboswitches, however, structural data might refer to mutually exclusive bound and unbound states. In this case, while feeding structures into the model, it may be more beneficial to treat these sequence-structure pairs separately as opposed to having them belonging to the same sequence. In any case, further investigation into different classes and types of RNAs may shed more light on the performance of Geometric GNNs on RNAs and a possible need for class-specific architectures for RNA computational design.

## Notes

### Competing Interest Statement

The authors have declared no competing interest.

